# The Effect of Visual Capture Towards Subjective Embodiment Within the Full Body Illusion

**DOI:** 10.1101/397943

**Authors:** Mark Carey, Laura Crucianelli, Catherine Preston, Aikaterini Fotopoulou

## Abstract

Typically, multisensory illusion paradigms emphasise the importance of synchronous visuotactile integration to induce subjective embodiment towards another body. However, the extent to which embodiment is due to the ‘visual capture’ of congruent visuoproprioceptive information alone remains unclear. Thus, across two experiments (total *N* = 80), we investigated how mere visual observation of a mannequin body, viewed from a first-person perspective, influenced subjective embodiment independently from concomitant visuotactile integration. Moreover, we investigated whether slow, affective touch on participants’ own, unseen body (without concomitant touch on the seen mannequin) disrupted visual capture effects to a greater degree than fast, non-affective touch. In total, 40% of participants experienced subjective embodiment towards the mannequin body following mere visual observation, and this effect was significantly higher than conditions which included touch to participants own, unseen body. The velocity of the touch that participants received (affective/non-affective) did not differ in modulating visual capture effects. Furthermore, the effects of visual capture and perceived pleasantness of touch was not modulated by subthreshold eating disorder psychopathology. Overall, this study suggests that congruent visuoproprioceptive cues can be sufficient to induce subjective embodiment of a whole body, in the absence of visuotactile integration and beyond mere confabulatory responses.

## 1. Introduction

Body ownership, the feeling that our body belongs to us and is distinct from other people’s bodies, is a fundamental component of our sense of self ^1,2^. Intuitively, this feeling appears stable and durable amongst humans, yet scientific studies have demonstrated that the sense of body ownership is a fragile outcome of integrating multiple sensory inputs. Such signals originate outside the body (i.e. exteroceptive modalities such as vision, touch) ^3,4^ as well as within the body (i.e. interoceptive modalities such as proprioception, heart rate) ^5–7^. These bodily signals are integrated to create a coherent sense of body ownership through which we interact with our environment ^2^.

Experimental paradigms have been successfully used to investigate how body ownership is shaped by the integration of incoming multisensory information. For example, in the *Rubber Hand Illusion (RHI)*^8^, individuals experience ownership over a fake (rubber) hand when placed in a congruent anatomical position and stroked in temporal synchrony with their own hand, which is hidden from view. This has been recently extended to ownership over an entire body *(Full Body Illusion)*, using a virtual avatar ^9^, or a mannequin body ^10^, viewed from a first-person perspective. In such illusions, the source of tactile stimulation on one’s own, unseen body (part) is attributed to the location of the visually perceived fake body (part) when the two are stroked synchronously, which is argued to give rise to subjective self-reports of illusory body ownership and a mislocation in one’s own sense of body position (i.e. proprioceptive drift) ^4^. Importantly, such effects typically occur within the constraints of top-down contextual factors, including the orientation ^3,11^, visual perspective ^12–14^, and appearance ^10,15,16^ of the embodied body (part). Indeed, research has shown that the strength of the illusion is modulated by the distance between the real and fake body (part), with greater spatial discrepancies decreasing the likelihood of integration between visuoproprioceptive signals ^17–19^.

Importantly, it has long been argued that the synchrony of the perceived touch with vision is a necessary condition for illusory ownership to occur, rather than asynchrony which is typically used as a control condition within multisensory illusion paradigms^11^. However, the role of synchronous visuotactile integration as a necessary component to trigger illusory embodiment remains debated ^20,21^. Research has shown that illusory embodiment could still be induced based purely on visual information of a fake body (part) in the absence of visuotactile stimulation ^21–23^, or based on merely expected but not experienced synchronous tactile stimulation ^24^, and even following asynchronous visuo-tactile stimulation, provided that spatial congruence is adhered to between the real and fake body (part) ^25^ (see ^20^ for review). Such evidence highlights that synchronous visuotactile input can strengthen illusory embodiment, by contributing to the downregulation in the weighting of proprioceptive signals regarding one’s own limb position in relation to vision ^26^, but may not be a necessary component to trigger this process ^21,22,27^.

However, studies which have investigated illusory body ownership in the absence of tactile stimulation have predominantly investigated this effect during the RHI (e.g. ^21,28,29^), with little research conducted towards a whole body ^12^. Among the latter, some have argued that synchronous visuotactile integration is a necessary condition to elicit illusory ownership in the full body illusion ^10^, while studies using virtual reality have found evidence to the contrary, following illusory ownership towards a virtual body in the absence of visuotactile integration ^9,12^. Therefore, we wished to investigate whether subjective visual capture of embodiment could occur towards a real mannequin body with a static field of view, from a first-person visual perspective in the ‘physical world’. In this context, ‘visual capture’ is defined as the degree of embodiment due solely to passive, visual perception of the fake body (part) viewed from a first-person perspective, independent from tactile stimulation (hereafter referred to as ‘visual capture of embodiment’) ^30,31^.

Interestingly, a tendency to weight visual information over other somatosensory signals has been recently observed in neuropsychological, right hemisphere patients with body representation deficits (e.g. ^31–34^). Moreover, clinical eating disorder patients have shown alterations in their weighting and integration of sensory information, which is argued to reflect an instability in the bodily self within this population ^35^. However, whilst ‘pure’ visual capture conditions have been tested in right hemisphere patients, evidence for heightened visual dominance within eating disorder patients derives from multisensory illusion studies finding that both synchronous and asynchronous visuotactile stimulation led to alterations in an individual’s body representation ^36–39^. Thus, direct investigation of visual capture of embodiment from congruent visuoproprioceptive cues alone (i.e. in the absence of tactile stimulation) has been less studied with regard to eating disorder psychopathology.

Importantly, greater illusory embodiment in acute eating disorder patients has been shown to persist to some degree amongst recovered patients, suggesting that such heightened sensitivity to visual information pertaining to the body may be a trait phenomenon ^37^. Therefore, such visual dominance over other sensory information may be independent from a status of malnutrition, and may occur *prior* to illness onset which could influence an individual’s body perception and body satisfaction ^40–42^. Thus, it may be that healthy individuals who display an increased visual capture of embodiment towards a fake body (part) show an increased visual dominance over other sensory information, which may link with a greater risk of developing distortions in body image. Consequently, the present study aimed to investigate whether subthreshold eating disorder psychopathology and body concerns may modulate the subjective embodiment shown towards a fake body as a result of mere visual capture.

In addition to research investigating visuoproprioceptive integration, the importance of interoception in multisensory integration and body ownership has only recently been investigated^7,43,44^. Interoception refers to information about the internal states of the body, processing sensations from within the body (e.g. hunger, thirst), but also outside the body (e.g. pleasure, pain), which is conveyed by a particular afferent pathway ^6^. Affective touch - i.e. slow, caress-like touch – is associated with increased pleasantness and has been found to activate specific C-Tactile (CT) afferents found only in the hairy skin, responding maximally to stroking velocities between 1 and 10 cm/sec ^45^. Importantly, affective tactile stimulation appears to be dissociable from exteroceptive, discriminatory stimulation such as non-affective touch^46^. Such CT afferents are hypothesised to take a distinct pathway to the posterior insular cortex^47,48^, an area associated with the early convergence of interoceptive information with exteroceptive bodily signals^6,49,50^.

Increasing evidence has shown that the velocity of perceived touch during visuotactile integration plays an influential role within the sense of body ownership. Specifically, touch delivered at CT-optimal velocities has been shown to increase embodiment during the RHI paradigm compared with fast, non-affective touch ^30,51–53^, however, evidence of this effect in the full body illusion remains equivocal ^54^. Moreover, recent research has shown that individuals with anorexia nervosa (AN) display a reduced subjective pleasantness to touch, relative to healthy controls ^49^; however, it is yet to be investigated how eating disorder psychopathology may modulate the extent to which individuals show alterations in their experience of touch, or vice versa. Therefore, within our second experiment, individual differences in the perception of touch will be investigated in relation to subthreshold eating disorder psychopathology.

In addition to enhancement of embodiment via interoceptive signals, evidence from patient populations with chronic pain has shown how feelings of body ownership can be disturbed ^55,56^ (but see ^57^ for review). Changes in interoceptive information (e.g. increased limb temperature) has been shown to disrupt the feelings of embodiment by *decreasing* the strength of the effect within multisensory illusions ^58^. Therefore, in addition to mere visual capture towards subjective embodiment (*visual capture* condition), the present study aimed to investigate the effects of tactile stimulation administered to participants’ own, unseen arm during visual observation of the mannequin body, as a control condition designed to ‘disrupt’ visual capture by introducing sensory input that is incongruent with participants’ visual information (*tactile disruption* condition). Furthermore, we aimed to investigate whether CT-optimal, affective touch (i.e. touch administered in CT-optimal velocities) would provide additional interoceptive information on one’s own body which would be expected to disrupt visual capture of embodiment to a greater extent compared with discriminatory, non-affective touch. Previous research has suggested that the perception of interoceptive signals depends on an individual’s ability to regulate the balance between interoceptive and exteroceptive information in ambiguous contexts ^7,30,59^. Thus, differences in an individual’s sensitivity and balance between these two streams of information may determine the degree of embodiment change shown during tactile disruption conditions.

In brief, we investigated whether mere visual observation of a mannequin body would lead to subjective embodiment when visuoproprioceptive cues are congruent with one’s own body. Based on previous research ^12,21^, we predicted that a compatible first-person perspective of a mannequin body would be sufficient to elicit subjective embodiment amongst participants, independent of concomitant tactile stimulation. In addition, we investigated the extent to which subjective embodiment towards the mannequin body was reduced when visual capture of proprioception was disrupted by tactile stimulation to participant’s own, unseen arm. We manipulated the velocity of tactile stimulation that participants received, to investigate whether slow, affective touch had a differential effect on the disruption of embodiment compared with fast, non-affective touch. Specifically, we predicted that the increased interoceptive information associated with affective touch would disrupt the downregulation of proprioceptive signals by visual capture to a greater extent compared to non-affective touch. Finally, we investigated whether subthreshold eating disorder psychopathology modulated any individual differences in subjective embodiment from visual capture. We hypothesized that higher eating disorder vulnerability would be associated with an increased weighting of visual information, and thus increased visual capture of embodiment. The above measures were replicated across two experiments, with the addition of a separate touch task in Experiment 2, designed to investigate the role of subjective pleasantness of touch in relation to subthreshold eating disorder psychopathology. Extending upon findings from clinical populations ^49^, we expected to observe a negative relationship between the above two measures, such that individuals with higher eating disorder psychopathology were hypothesised to display a reduced pleasantness to both affective touch and non-affective touch.

## 2. Methods

### 2.1 Experiment 1

#### 2.1.1 Participants

Forty-one healthy female participants (Mean age = 20.10, SD ± 2.48, range = 18-31) were recruited via the University of York research participation scheme and received course credit for a single 60-minute testing session. All participants had a healthy BMI (Mean = 21.48, SD ± 2.40, range = 18.30-28.60), no current or previous neurological or psychological disorders (self-report), and normal or corrected-to-normal vision. Exclusion criteria included any specific skin conditions (e.g. eczema, psoriasis) or any scarring or tattoos on the left arm. All participants gave informed consent to take part in the study. The study received ethical approval from the University of York Departmental Ethics Committee and was conducted in accordance with the Declaration of Helsinki. One participant was later excluded because she self-reported a previous psychological condition, therefore, the final sample consisted of forty participants (Mean age = 20.15, SD ± 2.49, range = 18-31).

#### 2.1.2 Design

The experiment employed a within-subjects design to investigate the effects of visual and tactile signals towards the subjective embodiment of a mannequin body. First, during *visual capture* trials participants visually observed the mannequin body for 30 seconds, from a first-person perspective, independent of any tactile stimulation. Second, participants also undertook trials identical to the *visual capture* condition, but with the addition of tactile stimulation applied (only) to participant’s own, unseen arm, designed to disrupt such visual capture (*tactile disruption* condition) for 60 seconds. Stimulation was administered at two different velocities to give rise to affective (3cm/s) and non-affective (18 cm/s) *tactile disruption*. The dependent variable was the subjective embodiment experienced by participants, rated after each trial via an *embodiment questionnaire* (see *Measures* section and Table 1 for details). The same *embodiment questionnaire* was completed for both *visual capture* and *tactile disruption* conditions. Participants completed two *visual capture* trials, each followed by an affective or non-affective *tactile disruption* trial in counterbalanced order between participants, resulting in a total of 4 trials per participant (see Figure 1).

**Table 1.** Embodiment Questionnaire presented to participants following each trial.

| Questionnaire Statement | Component |
| --- | --- |
| 1. It seemed like I was looking directly at my own body, rather than a mannequin body | Ownership |
| 2. It seemed like the mannequin body belonged to me | Ownership |
| 3. It seemed like the mannequin body was part of my body | Ownership |
| 4. It seemed like the mannequin body was in the location where my body was. | Location |
| 5. It felt like I had two bodies (at the same time) | Control |
| 6. It felt like my body was made out of rubber | Control |

**NB.** The order of questionnaire statements was randomized for each trial and participant.

**Figure 1.**
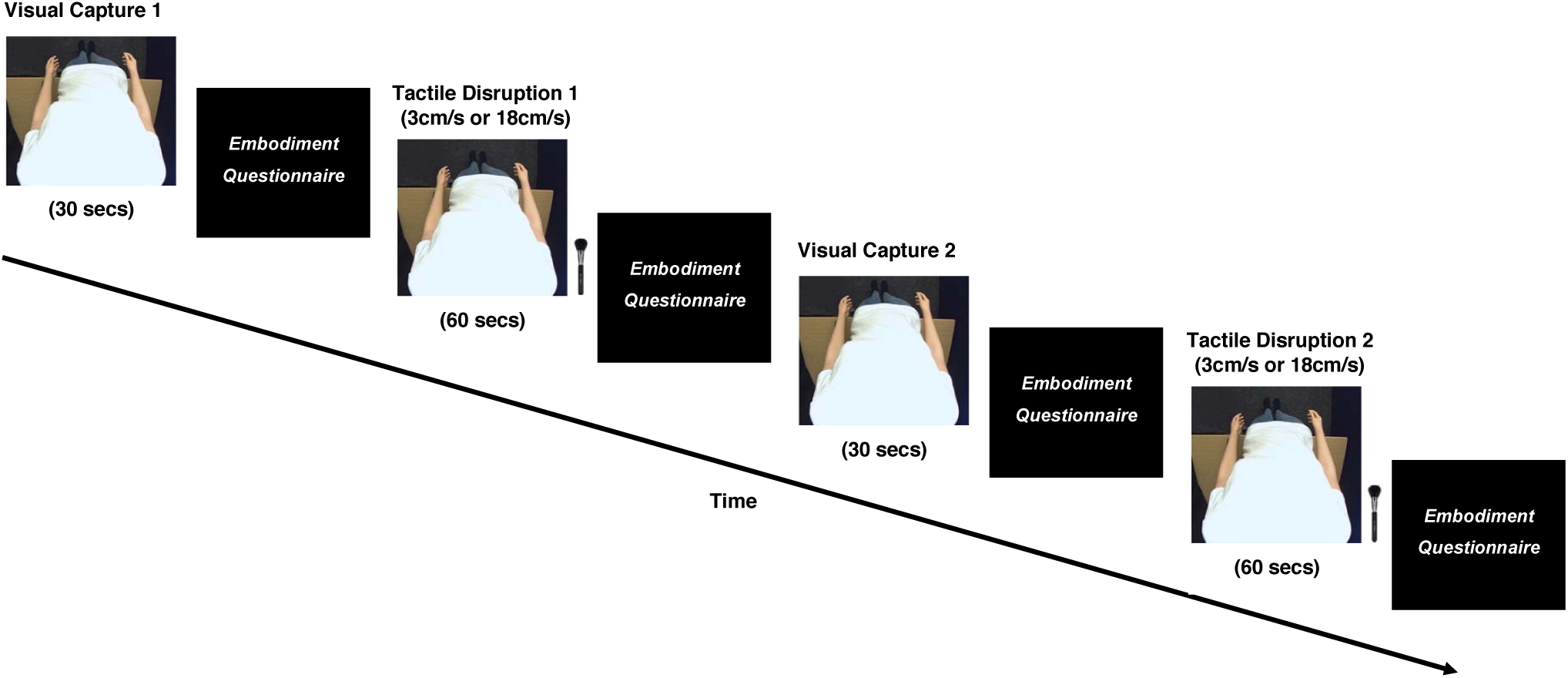
Timeline of experimental procedure. Participants completed two *visual capture* (30 secs) conditions and two *tactile disruption* (60 secs) conditions (1x affective touch; 1x non-affective touch). Tactile disruption order was counterbalanced across participants. Participants removed the HMDs following each trial and completed the *Embodiment Questionnaire* on a separate computer.

#### 2.1.3 Measures

##### 2.1.3.1 Embodiment Questionnaire

Following each trial, participants rated their subjective embodiment via an *embodiment questionnaire* (see Table 1) along a 7-point Likert scale (−3 strongly disagree to +3 strongly agree). This questionnaire (adapted from Longo et al., 2008) was composed of two subcomponents: *ownership* (i.e. the feeling that the mannequin body belongs to them) and *location* (i.e. the feeling that the mannequin body was in the position of their own body). An overall *embodiment score* was calculated by averaging the above two subcomponent scores. The final two statements were used as control items, in which an overall *control score* was calculated by averaging across the two control items. This was used within the *visual capture* condition to compare with *embodiment* scores, to ensure that responses from participants were specific to subjective embodiment and not due to task compliance.

##### 2.1.3.2 Eating Disorder Examination Questionnaire (EDE-Q) 6.0

The EDE-Q is a 28-item questionnaire used as a self-report measure of eating disorder psychopathology^60^ amongst community populations. The questionnaire assesses frequency of disordered eating behaviours (6 items), as well as eating behaviours and attitudes (22 items) within the past 28 days, along four subscales: *Dietary Restraint, Eating Concern, Weight Concern* and *Shape Concern*, which are also averaged for a *Global EDE-Q Score*. Items are rated along a 7-point (0-6) Likert scale, with higher scores signifying greater eating disorder psychopathology. This measure has good internal consistency, with Cronbach’s alpha ranging from .78 to .93 in a non-clinical sample ^61^. The overall global EDE-Q measure in the present study had a Cronbach’s alpha of .95 in both Experiment 1 and Experiment 2.

#### 2.1.4 Materials

A life-size female mannequin was used within the experimental set-up. The mannequin was dressed in a white t-shirt, blue jeans and black socks, with the head removed at the neckline to enable correct positioning of the video cameras. The body had a waist circumference of 62cm and was in a standing position with arms placed by their side (see Figure 2). During all trials, participants wore a set of head-mounted displays (HMDs) (Oculus Rift DK2, Oculus VR, Irvine, CA, USA), with a resolution of 1200 × 1080 pixels per eye, a refresh rate of 75Hz, and a corresponding nominal visual field of 100°. The HMDs were connected to a stereoscopic camera (USB 3.0 VR stereo camera, Ovrvision Pro, Japan), presenting a real time, video image to participants. The cameras were mounted and positioned downwards, at the eye line of the mannequin, capturing a first-person perspective of the body, compatible with looking down towards one’s own body. During *tactile disruption* trials, tactile stimulation was applied using a cosmetic make-up brush (Natural hair Blush Brush, N°7, The Boots Company). All experimental trials and responses were made using PsychoPy 2 ^62^ on an Apple iMac desktop computer (1.6GHz dual-core Intel Core i5 processor).

**Figure 2.**
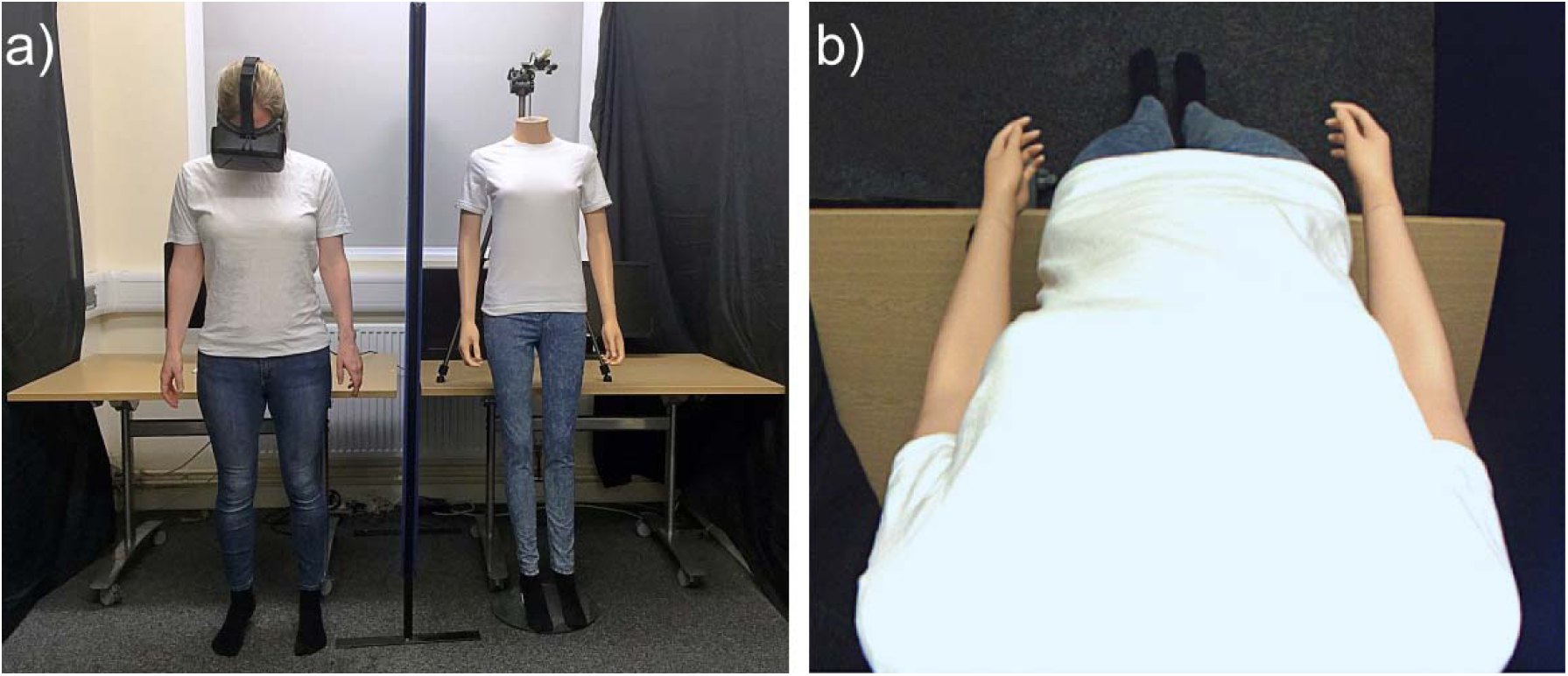
Experimental set-up. a) In *visual capture* trials, participants stood in an identical stance to the mannequin body, separated by a screen divider. b) Participants viewed a live video image of the mannequin from a first-person perspective, via head mounted displays.

#### 2.1.5 Experimental Procedure

Prior to the experiment, two adjacent 9×4cm stroking areas were marked on the hairy skin of each participants’ left forearm, using a washable marker pen ^51,63^. This provided a specific area for which to administer tactile stimulation for participants. Stimulation alternated between these two stroking areas within each *tactile disruption* trial, to minimise habituation, and provide the experimenter with an assigned area to control the pressure of each stroke. For all experimental trials, participants stood to the right of the mannequin body, separated by an office screen divider (see Figure 2a), whilst wearing the HMDs. Participants were instructed to remain still, place their arms by their side, and look down as though towards their own body. A live video image of the mannequin body, viewed from a first-person perspective, appeared in place of their own body through the HMDs (see Figure 2b).

For *visual capture* trials, participants visually observed the mannequin body for a 30-second period, without any tactile stimulation. Immediately after the trial, participants removed the HMDs and rated their subjective embodiment towards the mannequin via the *embodiment questionnaire* (see Table 1) on a separate computer. Removing the HMDs following each trial also served as a rest period for participants to move freely and dissociate their subjective experience between trials. For *tactile disruption* trials, participants identically visually observed the mannequin body, with the experimenter stroking participants’ own, unseen arm for a 60-second period. Stroking velocity was manipulated by administering slow, affective touch (3cm/s), and fast, non-affective touch (18cm/s). The experimenter was trained to administer each stroke at the precise speed, by counting the number of strokes within a window of 3 seconds per individual stimulation (i.e. one 3s-long stroke for 3 cm/s velocity, and six 0.5s-long strokes for 18 cm/s velocity). Identically, immediately after *tactile disruption* trials, participants removed the HMDs and rated their subjective embodiment towards the mannequin via the *embodiment questionnaire*. Individual questionnaire items were presented in a randomized order across all trials.

### 2.2 Experiment 2

#### 2.2.1 Participants

Forty-three healthy female participants (Mean age = 18.98, SD ± .74, range = 18 - 20) were recruited via the University of York research participation scheme and received course credit for a single 60-minute testing session. As in Experiment 1, all participants had a healthy BMI (Mean = 21.89, SD ± 2.67, range = 16.66-28.32), no current or previous neurological or psychological disorders (self-report), and normal or corrected-to-normal vision. Exclusion criteria included any specific skin conditions (e.g. eczema, psoriasis) or any scarring or tattoos on the left arm. All participants gave informed consent to take part in the study. The study received ethical approval from the University of York Departmental Ethics Committee and was conducted in accordance with the Declaration of Helsinki. Three participants were later excluded; one following a self-reported previous psychological condition; one excluded with scarring on their arms, and one excluded following poor comprehension with the experimental procedure. Therefore, the final sample consisted of forty participants (Mean age = 18.98, SD ± .77, range = 18 - 20).

#### 2.2.2 Design, Materials, Measures, Procedure

Design, Materials, Measures and Procedures were identical to Experiment 1, with the addition of a separate *Touch Task* completed prior to the *Full Body Illusion*, which explored subjective pleasantness ratings of affective vs. non-affective touch based solely on tactile input, in relation to subthreshold eating disorder psychopathology amongst healthy females.

##### Touch Task

Participants were asked to place their left arm on the table with their palm facing down, and wore a blindfold over their eyes to prevent any visual feedback to tactile stimulation. Tactile stimulation was administered using an identical cosmetic make-up brush (see *Materials* above) for 3 seconds per trial, at the same velocities as those in the *tactile disruption* conditions (affective touch - 3 cm/sec and non-affective touch - 18 cm/sec). There was a total of six trials per velocity condition, for a total of 18 trials, with all trials presented in a randomized order for each participant. Following each trial, participants verbally reported the pleasant of the touch, using the pleasantness rating VAS scale, anchored from 0 (*Not at all pleasant)* to 100 (*Extremely pleasant*) ^49^. An average score across the six trials was calculated to obtain a single score, per participant, for each of the two tactile conditions.

### 2.3 Data Analysis

All statistical analyses were conducted using SPSS version 23.0 (IBM, Chicago, IL, USA). Data from the *embodiment questionnaire* were ordinal and found to be non-normal via a Shapiro-Wilk test (*p* < .05), thus, appropriate non-parametric tests were used for analysis. Data for pleasantness ratings in the *Touch Task* were normally distributed (*p* > .05), therefore parametric tests were used to analyse this data. Effect sizes for parametric tests are indicated by Cohen’s *d*, and non-parametric Wilcoxon signed-rank tests are indicated by r values (*r*) which are equivalent to Cohen’s *d* ^64^.

To examine whether mere visual observation of a mannequin body would lead to subjective embodiment (*visual capture* effect) we used a Wilcoxon signed-rank test to compare *embodiment* scores with *control* scores within the *embodiment questionnaire* (see Table 1 for *embodiment questionnaire* items). In addition, to investigate whether subjective embodiment was significantly reduced when visual capture was disrupted by tactile stimulation to participant’s own, unseen arm (*tactile disruption*), a further Wilcoxon signed-rank test was conducted to compare subjective *embodiment* scores between *visual capture* and *tactile disruption* conditions. Moreover, we assessed whether slow, affective touch on participants own arm led to greater disruption in subjective embodiment within participants compared with fast, non-affective touch, using a Wilcoxon signed-rank test to compare *embodiment* scores between the two stroking velocities (affective vs. non-affective *tactile disruption*). The above analyses were also conducted for individual *Ownership* and *Location* subcomponents within the *embodiment questionnaire* (see Supplementary Materials, Sections 1 and 2). In addition, in Experiment 2 we examined the effect of stroking velocity on pleasantness ratings using a paired-samples t-test, to first establish whether slow, affective touch was indeed perceived as significantly more pleasant that fast, non-affective touch (manipulation check). The perception of touch was then investigated in relation to subthreshold eating disorder psychopathology (as measured by the EDE-Q 6.0), using a non-parametric Spearman’s correlation.

To establish individual differences in reported visual capture of embodiment, we calculated percentage frequencies across the combined samples of Experiment 1 and 2, of those who reported visual capture of embodiment (average scores of ≥ +1 in response to the *embodiment questionnaire* ^3,65^), those who neither affirmed or denied embodiment (average scores of < +1 and > −1 in response to the *embodiment questionnaire*) and those who denied visual capture (average scores of < −1 in the *embodiment questionnaire*). Finally, we wished to explore whether such individual differences in subjective embodiment from visual capture related to subthreshold eating disorder psychopathology (EDE-Q 6.0). Therefore, we conducted a non-parametric Spearman’s correlational analysis between the psychometric EDE-Q measure and subjective embodiment scores from *visual capture*.

## 3 Results

### 3.1 Experiment 1

#### 3.1.1 Embodiment Questionnaire

Preliminary analysis showed that there was no effect of trial order across visual capture trials, with a Wilcoxon signed-rank test revealing no significant difference in embodiment scores between visual capture trial 1 vs. trial 2 (*Z* = - .084, *p* = .933). Therefore, *embodiment questionnaire* scores were collapsed across the two visual capture trials to provide an overall *visual capture* embodiment score, per participant.

##### 3.1.1.1 Main effect: Visual Capture

To examine the effects of mere visual capture towards subjective embodiment of the mannequin body, we compared *embodiment* scores with *control* scores in the *embodiment questionnaire*. A Wilcoxon signed-rank test revealed a main effect of visual capture, with significantly higher embodiment scores compared with control scores (*Z* = −4.04, *p* < .001, *r* = 64) (see Figure 3).

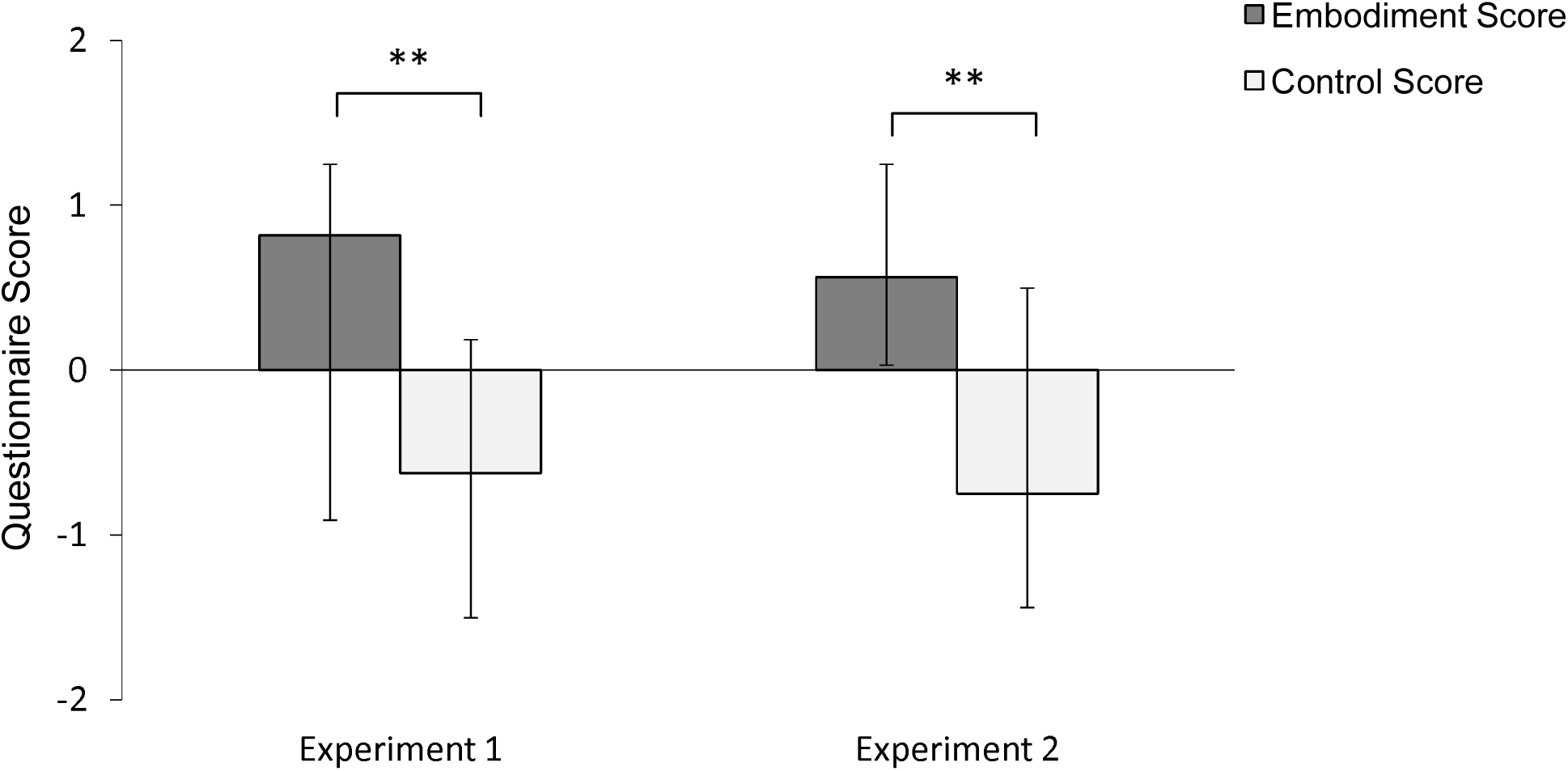

##### 3.1.1.2 Main effect: Tactile Disruption

In order to determine whether tactile disruption to participants’ own unseen arm would disrupt subjective embodiment, we compared *embodiment* scores between *tactile disruption* and *visual capture* conditions. A Wilcoxon signed-rank test revealed a main effect of condition, in which participants showed significantly lower subjective embodiment following *tactile disruption* trials (median = −.38) compared with *visual capture* trials (median = .82) (Z = −3.74, *p* < .001, *r* = .59).

##### 3.1.1.3 Main effect: Tactile Velocity

Next, we examined whether tactile velocity had an effect in disrupting the subjective embodiment towards the mannequin body within *tactile disruption* trials. A Wilcoxon signed-rank test revealed that there was no significant difference in embodiment scores between affective and non-affective tactile disruption trials (*Z* = −.104, *p* = .918, *r* = .02), which suggests that interoceptive affective touch did not disrupt visual capture of embodiment to a greater degree than exteroceptive, non-affective touch.

### 3.2 Experiment 2

#### 3.2.1 Touch Task (Manipulation Check)

A further one participant was later excluded within the *Touch Task* analysis as an extreme outlier, scoring more than 2 SD below the group mean in pleasantness ratings of affective touch (3cm/s velocity) ^29^. Therefore, the final sample for this analysis consisted of 39 participants. As expected, a paired samples t-test revealed an effect of stroking velocity within the touch task, with significantly higher subjective pleasantness ratings following affective touch (3cm/s) (mean = 74.27) compared with non-affective touch (18cm/s) (mean = 52.94) (*t* (38) = 7.93, *p* < .001, *d =* 1.27). Moreover, correlational analyses were conducted to investigate the relationship between pleasantness ratings and subthreshold eating disorder psychopathology (measured by the *Eating Disorder Examination Questionnaire*; EDE-Q 6.0). First, a Spearman’s rank correlation revealed an approaching significant correlation between pleasantness ratings (average affective/non-affective touch) and global EDE-Q score (*r* = −.316, *p* = .05). Next, difference scores were calculated between affective and non-affective touch pleasantness ratings to determine whether those with higher subthreshold eating disorder psychopathology were less sensitive to differences in the affectivity of touch. However, a Spearman’s rank correlation revealed no significant correlation between touch difference score and global EDE-Q (*r* = .014, *p* = .935). Thus, the results suggest a trend in which those scoring higher in subthreshold eating disorder psychopathology may show a reduced pleasantness to all tactile stimulation, however this may not be further modulated by the affectivity of the touch that they receive.

#### 3.2.2 Embodiment Questionnaire

Preliminary analysis showed that there was no effect of trial order across visual capture trials, with a Wilcoxon signed-rank test revealing no significant difference in embodiment scores between visual capture trial 1 vs. trial 2 (*Z* = - .958, *p* = .338). Therefore, *embodiment questionnaire* scores were collapsed across the two visual capture trials to provide an overall *visual capture* embodiment score, per participant.

##### 3.2.2.1 Main effect: Visual Capture

To examine the effects of mere visual capture towards subjective embodiment of the mannequin body, we compared *embodiment* scores with *control* scores in the *embodiment questionnaire*. A Wilcoxon signed-rank test revealed a main effect of visual capture, with significantly higher embodiment scores compared with control scores (*Z* = −4.30, *p* < .001, *r* = .68) (see Figure 3).

##### 3.2.2.2 Main effect: Tactile Disruption

In order to determine whether tactile disruption to participants’ own unseen arm would disrupt subjective embodiment, we compared embodiment scores between *tactile disruption* and *visual capture* conditions. A Wilcoxon signed-rank test revealed a main effect of condition, in which participants showed significantly lower subjective embodiment following tactile disruption trials (median = −.23) compared with visual capture trials (median = .59) (Z = −4.08, *p* < .001, *r* = .65).

##### 3.2.2.3 Main effect: Tactile Velocity

Next, we examined whether tactile velocity had an effect in disrupting the subjective embodiment towards the mannequin body within *tactile disruption* trials. A Wilcoxon signed-rank test revealed that there was no significant difference in embodiment scores between affective and non-affective tactile disruption trials (*Z* = - .354, *p* = .723, *r* = .06), which suggests that interoceptive affective touch did not disrupt embodiment to a greater degree than exteroceptive, non-affective touch.

### 3.3 Combined Samples

#### 3.3.1 Visual Capture of Embodiment – Individual Differences

Across the combined, total sample (*N*=80), 32 participants (40%) experienced a degree of embodiment over the mannequin from mere visual capture, with average scores of ≥ +1 in response to the *embodiment questionnaire* (hereafter referred to as ‘visual capture’ (VC) group). To confirm this percentage was not a consequence of participant compliance, a Wilcoxon signed rank test was conducted which revealed a significant difference between *embodiment* and *control* scores (Z = −4.71, *p* < .001, *r* = .74), with only 4 participants (12.5%) of the VC group scoring ≥ +1 in response to *control* items. 36 participants (45%) seemed to neither affirm or deny embodiment over the mannequin, with average scores of < +1 and > −1 in response to the *embodiment questionnaire* (hereafter referred to as ‘borderline’ group). 12 participants (15%) of the total sample denied any subjective embodiment from visual capture, with average scores of < −1 in the *embodiment questionnaire* (hereafter referred to as ‘no visual capture’ (no-VC) group).

#### 3.3.3 Subthreshold Eating Disorder Psychopathology

Finally, correlational analyses were conducted to investigate the relationship between visual capture effects and subthreshold eating disorder psychopathology (measured by the *EDE-Q 6.0*). EDE-Q subscale and global scores across both experiments are presented in Table 2. A Spearman’s rank correlation revealed no significant correlation between visual capture embodiment scores and global EDE-Q scores (*r* = .030, *p* = .79), or any EDE-Q subscale scores (all *ps* > .05). Similarly, no significant correlations were observed when analysing subcomponent (*Ownership* and *Location*) scores within the *embodiment questionnaire* with EDE-Q scores (see Supplementary Materials, Section 3). This suggests that subthreshold attitudes and behaviours regarding to eating and body image did not relate to the degree of subjective embodiment of a mannequin body due to mere visual capture.

**Table 2.** Participant demographic information (Mean and (SD)) and EDE-Q subscale and global scores.

|  | Total (N=80) | Experiment 1<br>(N=40) | Experiment 2<br>(N=40) | *t* | *p* |
| --- | --- | --- | --- | --- | --- |
| Age | 19.56 (1.92) | 20.15 (2.49) | 18.98 (.77) | 2.86 | .006 |
| BMI | 21.70 (2.56) | 21.48 (2.40) | 21.93 (2.71) | -.772 | .442 |
| Restraint | .80 (.20-1.80)<sup>a</sup> | .80 (.20-2.15)<sup>a</sup> | .90 (.25-1.75)<sup>a</sup> | -.101<sup>b</sup> | .919 |
| Eating Concern | .60 (.20-1.40)<sup>a</sup> | .60 (.20-1.40)<sup>a</sup> | .60 (.20-1.55)<sup>a</sup> | -.567<sup>b</sup> | .571 |
| Shape Concern | 2.25 (1.16-3.72)<sup>a</sup> | 2.06 (1.25-3.63)<sup>a</sup> | 2.31 (1.00-3.75)<sup>a</sup> | -.106<sup>b</sup> | .916 |
| Weight Concern | 1.40 (.40-3.00)<sup>a</sup> | 1.40 (.40-2.55)<sup>a</sup> | 1.70 (.50-3.20)<sup>a</sup> | -.960<sup>b</sup> | .337 |
| EDE-Q Global | 1.33 (.60 -2.32)<sup>a</sup> | 1.31 (.60-2.17)<sup>a</sup> | 1.35 (.65-2.52)<sup>a</sup> | -.380<sup>b</sup> | .704 |

<sup>a</sup>Median and interquartile range in parentheses
<sup>b</sup>Mann-Whitney U statistic

## Data Availability

The datasets analysed during the current study are available from the corresponding author on reasonable request.

## 4. Discussion

The present study investigated the extent to which mere visual observation of a mannequin body, viewed from a first-person perspective, influenced subjective embodiment independently from concomitant visuotactile integration. Across two experiments, our results showed that congruent visuoproprioceptive cues between one’s own physical body and a mannequin body was sufficient to induce subjective embodiment in 40% of our total sample. Furthermore, as expected, embodiment was significantly reduced when ‘visual capture’ of embodiment was disrupted by tactile stimulation to participant’s own, unseen arm, confirming that the visual capture effect on embodiment was not due to confabulatory or social desirability responses. Contrary to our secondary hypothesis regarding interoception, this tactile disruption effect was not modulated by stroking velocity, with comparable changes in embodiment following slow, affective (CT-optimal) and fast, non-affective touch. Finally, subthreshold eating disorder psychopathology was not found to modulate the effects of embodiment in visual capture or tactile disruption conditions.

Our findings support previous research which argues that synchronous visuotactile stimulation is not a necessary condition amongst all individuals in triggering subjective embodiment within bodily illusions. Research has shown that visual capture of proprioception can be sufficient to elicit embodiment towards a fake hand ^21,31^ and whole body ^12^ in some individuals. Indeed, whilst Maselli and Slater (2013) have shown this effect using a full body within an immersive, virtual environment, the present study is the first to explore this effect towards a full body in the ‘physical world’. Our results suggest that multisensory illusion paradigms would benefit from a baseline measure based on the mere visual observation of the fake body (part) (i.e. visual capture effect), which is unbiased by concomitant visuotactile stimulation ^30,51^. Indeed, this is in support of research which argues that asynchronous stimulation in multisensory illusion paradigms is not strictly a neutral, control condition within multisensory body illusions ^25,28^, with visuotactile asynchrony instead providing somatosensory conflict ^25,66^.

The present data showed that a substantial percentage of participants displayed a degree of subjective embodiment towards the mannequin body following mere visual observation. Indeed, it was confirmed that such individuals who did display visual capture of embodiment were not simply complying with all items in the *embodiment questionnaire,* shown by significantly higher responses in *embodiment* scores compared with *control* scores (see *Results* section). However, congruent visuoproprioceptive signals did not induce subjective embodiment amongst all individuals to the same degree. We speculate that such individual differences may be due to a number of processes; for example, some individuals may have weaker proprioceptive signals which would give rise to greater sensory weighting towards the salient visual cues of the mannequin body within the illusion. Indeed, our own hypothesis that individual differences in visual capture may relate to subthreshold eating disorder psychopathology was not confirmed (see below for further discussion). Thus, further research is required to establish how individual differences in the weighting of distinct sensory cues contribute to modulating body ownership in mere visual capture conditions.

Furthermore, our results showed that tactile stimulation to participants own, unseen arm significantly disrupted subjective embodiment towards the mannequin body, by delivering somatosensory information that was incongruent with participants visuoproprioceptive cues. This result further highlights that the embodiment shown from visual capture conditions were not due to participant compliance, as disruption to such visual capture resulted in significantly lower embodiment scores. From a computational approach to multisensory integration ^21,26,67^, such incongruent tactile information is likely to have disrupted the sensory weighting that is occurring between visual and proprioceptive body signals. Indeed, predictive coding accounts of multisensory illusions argue that illusory embodiment typically occurs by the brain downregulating the precision of conflicting, bottom-up somatosensory signals, which allows top-down predictions to resolve any sensory ambiguity about the body (i.e. *the body (part) I see is mine*) ^26^. Therefore, in the present study, additional tactile input to participants’ own, unseen arm added further somatosensory information which could not be downregulated or “explained away” by top-down predictions, given its incongruency with the visually perceived mannequin body ^68^, thus leading to reduced subjective embodiment.

Moreover, it was expected that the interoceptive properties associated with slow, affective touch^30^ would disrupt subjective embodiment to a greater degree than fast, non-affective touch. This is following evidence that affective touch led to enhanced embodiment during RHI paradigms ^51–53^, which is argued to be due to the additional interoceptive information conveyed by this CT-optimal touch ^69^. Further, research has shown that manipulation of interoceptive information (e.g. changes in body temperature) can *disrupt* feelings of body ownership ^58^. However, contrary to our predictions and previous findings, our results showed that the interoceptive, affective tactile stimuli did not appear to disrupt visual capture of embodiment to a greater extent than non-affective tactile stimuli. Such findings may be because the salience of incongruent visuotactile information was sufficient in disrupting embodiment towards the mannequin, with the subtlety of increased interoceptive information from the arm following affective touch providing no additional value to multisensory integration in this context. Furthermore, the previously observed effects of affective touch in enhancing body ownership during the RHI (which involves concomitant felt and seen touch on the rubber hand) may also be explained by the vicarious affectivity of the *seen* touch in addition to the interoceptive nature of the felt touch (Filippetti et al., submitted). Indeed, CT-optimal velocities have been shown to have distinct vicarious touch effects in behavioural ^70^ and neuroimaging ^50^ studies. However, visual cues of affective touch were not present in the current study, therefore the felt affectivity of the touch may have been attenuated by participants receiving only tactile stimulation that was not visually observed.

The present results must be considered in relation to the top-down, cognitive constraints within which illusory ownership is argued to occur. Research has shown that the embodied fake body (part) must be in an anatomically plausible position ^3,11,18,19^, must represent a corporeal object ^10,15,16^, and must be viewed from a first-person visual perspective ^12–14^. Indeed, it has been shown that when these constraints are violated, illusory effects diminish or disappear ^20,71,72^, suggesting that the perceived fake body (part) is required to fit with a reference model of the body based on top-down information^16^. The above conditions were closely adhered to in the present study, which was particularly salient using the HMDs, allowing a high degree of spatial overlap by *replacing* the first-person perspective of one’s own body with the identical perspective of a mannequin body. This provided a greater congruence of visuoproprioceptive cues which cannot be as closely matched within the RHI set-up without the use of computer-generated technology. However, further research should investigate the specific boundaries within which mere visual capture is sufficient in inducing embodiment towards a whole body, in the absence of visuotactile stimulation^12,73^, by systematically manipulating the above conditions within which the illusion can typically occur.

Finally, following evidence that acute eating disorder patients display a dominance in weighting to visual information related to the body^36,38^, which is shown to persist after recovery^37^, we explored whether this trait phenomenon would exist amongst healthy individuals, in relation to subthreshold eating disorder symptomology. However, no significant correlations were observed between EDE-Q scores and subjective embodiment following *visual capture*. This finding is in line with previous research in which those higher in subthreshold eating disorder symptoms did not experience a stronger *subjective* embodiment within the full body illusion^42^, despite relationships observed between EDE-Q scores and subsequent behavioural measures (e.g. body satisfaction) following the illusion (see also ^39^ for similar effects in AN patients). This suggests that previous findings which highlight differences in subjective embodiment in relation to eating disorder psychopathology may be body part specific ^36,38,74^. Nevertheless, studying eating disorder characteristics within healthy individuals remains clinically important to identify factors associated with the development of eating disorders without the confounds of physical consequences of the disorder ^75,76^.

Taken together, the present findings are in accordance with previous research which highlights the dynamic mechanisms that lead to illusory body ownership ^12^. First, there exists a two-way interaction between visual information of the fake body (part) and proprioceptive information of one’s own body (part), which is combined to inform an estimate of an individual’s current spatial position. When the fake body (part) is in an anatomically plausible position with one’s own body, sensory information between competing visual and proprioceptive cues is weighted in favour of the salient visual information ^67,77^, which for many is sufficient to induce feelings of embodiment to occur *prior* to visuotactile integration ^12,21^. Subsequently, the addition of synchronous visuotactile information creates a three-way weighted interaction between vision, touch and proprioception, with the visually perceived touch processed in a common reference frame based on the visuoproprioceptive cues. The subsequent ‘visual capture’ of synchronous visuotactile stimulation acts to further weaken one’s own proprioceptive signals, which can lead to increased illusory embodiment ^20,72^. Thus, future studies which compare the two-way vs. three-way interaction between sensory inputs would be informative in quantifying the additive effect that visuotactile stimulation plays within such paradigms. This could also be used to further investigate individual differences in the susceptibility to integrate visuoproprioceptive information to a greater degree than the additional integration of tactile stimuli during the illusion.

In conclusion, the present study suggests that mere visual observation of a mannequin body, viewed from a first-person perspective, can elicit subjective embodiment amongst individuals. Congruent visuoproprioceptive cues between one’s own physical body (part) and a fake body (part) was shown to be sufficient to induce subjective embodiment in 40% of our total sample in the absence of concomitant visuotactile stimulation, which is typically used to induce illusory embodiment within multisensory illusion paradigms. In addition, tactile stimulation delivered to participants own, unseen arm acted to disrupt reported subjective embodiment, however, this was not influenced to a greater degree by slow, affective touch compared with fast, non-affective touch. This suggests that interoceptive information about one’s body does not have the potency of discriminatory tactile signals, when the integration of vision and proprioception need to be moderated by touch. Future studies should explore this possibility using other interoceptive modalities such as cardiac awareness, and further investigate the role of sensory weighting and integration in clinical eating disorder populations.

## Acknowledgements

We are grateful to Chloe Robinson for her help and feedback in piloting the present study. The study was supported by a European Research Council Starting Investigator Award [ERC-2012-STG GA313755] (to AF), and a University of York studentship (to MC). Funding for the time of AF and LC has been partially provided by the Fund for Psychoanalytic Research through the American Psychoanalytic Association.

## Author Contributions

MC, LC, CP and AF designed the experiment. MC performed data collection and analysed the data, under supervision of CP, LC and AF. MC drafted the manuscript, and CP, LC and AF provided critical revisions. All authors approved the manuscript before submission.

